# Capturing multiple-type interactions into practical predictors of type replacement following HPV vaccination

**DOI:** 10.1101/523472

**Authors:** Irene Man, Kari Auranen, Jacco Wallinga, Johannes A. Bogaards

**Affiliations:** Centre for Infectious Diseases Control, National Institute for Public Health and the Environment, The Netherlands; Department of Medical Statistics and Bioinformatics, Leiden University Medical Center, The Netherlands; Department of Mathematics and Statistics, and Department of Clinical Medicine, University of Turku, Finland; Department of Epidemiology and Biostatistics, Vrije Universiteit, Amsterdam UMC, The Netherlands

**Author notes:** Present address: RIVM, P.O. Box 1, 3720 BA Bilthoven, The Netherlands.

**Keywords:** pathogen type interactions, vaccination, prediction, type replacement, hazard ratio, odds ratio

## Abstract

Current HPV vaccines target a subset of the oncogenic human papillomavirus (HPV) types. If HPV types compete during infection, vaccination may trigger replacement by the non-targeted types. Existing approaches to assess the risk of type replacement have focussed on detecting competitive interactions between pairs of vaccine and non-vaccine types. However, methods to translate any inferred pairwise interactions into predictors of replacement have been lacking. In this paper, we develop practical predictors of type replacement in a multi-type setting, readily estimable from pre-vaccination longitudinal or cross-sectional prevalence data. The predictors we propose for replacement by individual non-targeted types take the form of weighted cross hazard ratios of acquisition versus clearance, or aggregate odds ratios of coinfection with the vaccine types. We elucidate how the hazard-based predictors incorporate potentially heterogeneous direct and indirect type interactions by appropriately weighting type-specific hazards and show when they are equivalent to the odds-based predictors. Additionally, pooling type-specific predictors proves to be useful for predicting increase in the overall non-vaccine-type prevalence. Using simulations, we demonstrate good performance of the predictors under different interaction structures. We discuss potential applications and limitations of the proposed methodology in predicting type replacement, as compared to existing approaches.

## 1 Introduction

Predicting the impact of vaccination against a pathogen can be challenging if the pathogen consists of many, potentially interacting (sub)types. When the vaccine targets only a subset of the pathogenic types, it is particularly relevant to evaluate the risk of replacement by the nontargeted types. In this paper, we expand existing methodology for predicting type replacement for multi-type pathogens with the focus on the human papillomavirus (HPV).

HPV is one of the most common oncogenic DNA viruses in humans. Persistent infection with HPV can cause cancer in various parts of the body [1]. In particular, twelve to fifteen HPV types are classified as high-risk or probable high-risk due to their association with cervical cancer [2]. Currently, three HPV vaccines are available, covering two, four, and nine HPV types. All three vaccines protect against HPV 16 and 18, two high-risk types that together account for approximately 70% of all cervical cancers in unvaccinated populations. HPV 31, 33, and 45, accounting for an additional 10-15% of cases [3], are among the cross-protective types of the bivalent and quadrivalent vaccines and are included in the nonavalent vaccine [4].

In countries where HPV vaccination has been implemented with high coverage, circulation of both the vaccine and cross-protective HPV types has decreased considerably [5, 6]. However, concerns have been raised that the ecological niche vacated by the targeted types could be taken over by the non-targeted high-risk HPV types [7, 8, 9]. Thus far, post-implementation surveillance data have been reassuring, as only sporadic increases in the prevalence of infection with some non-vaccine types have been observed without a clear signal of type replacement [4, 10, 11, 12]. While waiting for the long-term impact of HPV vaccination to become apparent, evaluation of the potential for type replacement remains crucial.

Whether HPV vaccination will trigger type replacement depends on the existence and strength of competitive interactions between HPV types. HPV types may compete during coinfection, by either diminishing each other’s opportunities to establish a productive infection or by promoting viral clearance (e.g. through activation of antigen-presenting cells) [13, 14, 15, 16, 17]. Due to such competitive mechanisms, the hazard of acquiring (or clearing) a given HPV type may be reduced (or increased, respectively) by infection with other types. In epidemiological terms, different type interactions can be conveniently quantified in terms of appropriately defined hazard ratios [18, 19, 20].

We have previously shown, in a simple model of one vaccine and one non-vaccine type, that the cross hazard ratio of acquisition versus clearance can be used to predict type replacement, provided that the two types interact symmetrically and there is no long-lasting cross-immunity [21]. The latter assumption seems plausible for HPV as even the existence of natural homologous immunity is still debated [22, 23]. Given Susceptible-Infected-Susceptible (SIS) transmission dynamics, the appropriate cross hazard ratio is, moreover, equivalent to an odds ratio of coinfection, and thus estimable from cross-sectional prevalence data [21]. Here, the odds ratio is defined as the odds of infection with one type in presence versus absence of coinfection with the other type.

With more than two interacting types, not only does prediction of replacement require inference of interactions between multiple types but also an adequate way of combining them. To predict how vaccination will affect the prevalence of a given non-vaccine type, one needs to take account of its direct interactions with the vaccine types as well as any indirect interactions via other non-vaccine types. Meaningful prediction should also incorporate possible heterogeneities in strength or direction (competition versus synergy) of type interactions.

Previous studies evaluating the potential of HPV type replacement have focussed on inferring interactions between pairs of vaccine and non-vaccine types [7, 24, 25, 26, 27]. In these studies, for each vaccine type, pairwise odds ratios have been compared to pooled odds ratios (pooled across the non-vaccine types) to identify likely candidates for type replacement. With this approach, however, each non-vaccine type is evaluated multiple times according to its interactions with different vaccine types, while the occurrence of type replacement is determined by all these interactions jointly. In addition, the pooled odds ratio, which has been interpreted as the tendency of the vaccine type in question to be involved in coinfection, lacks a clear interpretation for prediction of type replacement.

In this paper, we consider prediction of replacement following vaccination in a setting with an arbitrary number of interacting vaccine and non-vaccine types. We propose predictors of type-specific replacement, i.e. increase in the prevalence of individual non-vaccine types, and pooled predictors of increase in the overall non-vaccine-type prevalence. The predictors are initially defined in terms of steady-state hazards of type-specific acquisition and clearance. Using a mechanistic SIS model, we explain how the predictors relate to the underlying mode of type interactions and show under which interaction structures these hazard-based predictors can be estimated as odds ratios from cross-sectional prevalence data. Finally, we evaluate the performance of the proposed predictors by means of numerical simulations.

## 2 Material and methods

### 2.1 Prediction framework

We derive predictors of type replacement for a pathogen consisting of many potentially interacting types. The prediction method applies to any pathogen for which naturally acquired immunity is short-lived, so that the transmission dynamics of each type can be approximated by an SIS model. The predictors are constructed in terms of the following data collected at the pre-vaccination steady state:

- prevalence of each (co)infection state, estimable from cross-sectional data;
- type-specific hazards (per capita rates) of acquisition and clearance, estimable from longitudinal data.

Figure 1a shows the eight (co)infection states and the respective transitions for a pathogen with three types.

**Figure 1:**
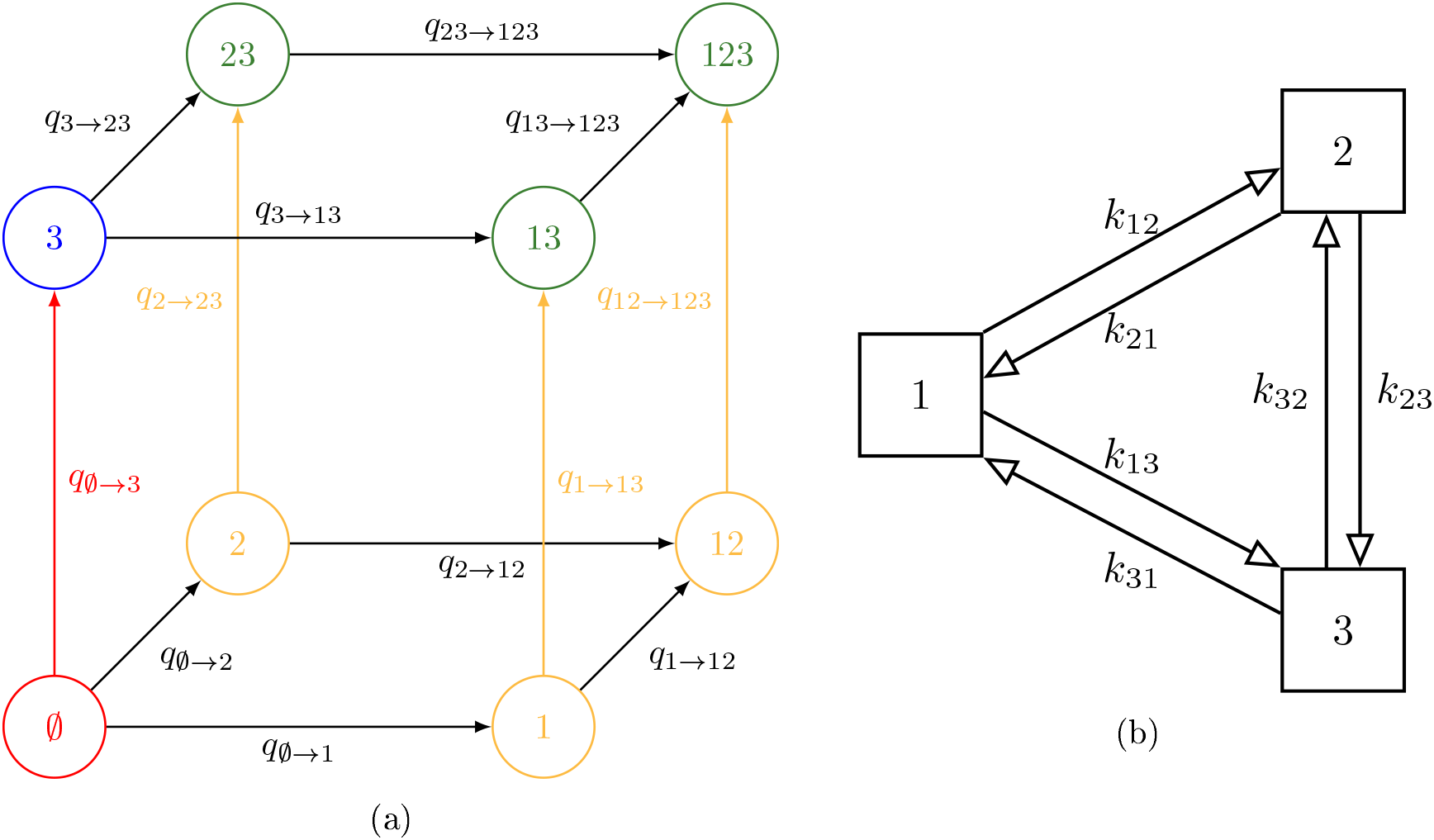
(a) The eight, infection states and transitions pertaining to acquisitions 111 a three-type system. For convenience, the reverse transitions (i.e. clearances) are not shown. The colours indicate the aggregate states for *VT* = {1, 2} *i* = 3 and the corresponding type-3 acquisitions; 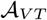 in yellow, 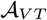 in green, 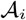 in blue, 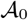 in red. (b) Graphical representation of the interaction parameters under the symmetric multiplicative structure or pairwise-asymmetric multiplicative structure 111 a three-type system. Under the pairwise-symmetric multiplicative structure, pairs of reverse interaction parameters would moreover be identical (i.e. *k_ij_* = *k_ji_*).

Type interaction is assumed to operate through current (co)infection, with one or multiple types modifying acquisition hazards of other types, or modifying clearance hazards of coinfecting types. Interactions between different types are allowed to be either competitive or synergistic and to vary in strength. Furthermore, we focus on predicting replacement by non-vaccine types that are not cross-protected by the vaccine, as these types are the most salient for evaluating the potential for replacement. Moreover, we ignore evolution through mutation of the model pathogen types for the timescale on which type replacement may occur.

### 2.2 Type-specific and overall replacement

We consider replacement, here defined as increase in the prevalence of non-vaccine-type infection once the vaccine types are eliminated in the post-vaccination steady state, at two levels:

- type-specific replacement, defined as increase in the prevalence of a given non-vaccine type *i*. This occurs when 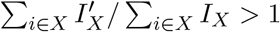, where 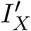 and *I_X_* denote the post- and prevaccination steady-state prevalence of infection state *X*, respectively. The sums here are taken over all states containing type *i*, e.g. states {3,13, 23,123} in a three-type system when considering *i* = 3;
- overall replacement, defined as increase in the overall non-vaccine-type prevalence. This occurs when 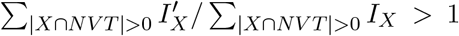 where *NVT* is the set of all nonvaccine types, and | · | denotes the number of types in a given set. The sums are taken over all states containing at least one non-vaccine type, e.g. states {2, 3, 23,12,13,123} in a three-type system when considering *NVT* = {2, 3}.

### 2.3 Predictors in a two-type setting

Previously, we have shown that the following pairwise odds ratio is an exact predictor of type replacement in a simple SIS model of one vaccine type (type 1) and one non-vaccine type (type 2), provided the two types interact symmetrically [21]:

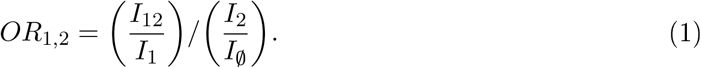

This pairwise odds ratio compares the odds of non-vaccine-type infection between those infected and uninfected with the vaccine type.

Whenever the pre-vaccination steady-state value of *OR*_1,2_ is less than one, replacement will occur. This correspondence follows from *OR*_1,2_ being equivalent to the following pairwise cross hazard ratio:

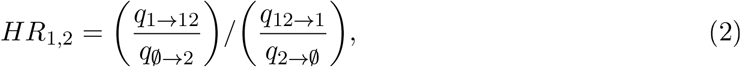

where *q_X→Y_* denotes the transition hazard from state *X* to *Y*. The numerator (denominator) of expression (2) is the hazard ratio of acquiring (clearing) the non-vaccine type in those infected versus those uninfected with the vaccine type and thus quantifies interaction in acquisition (clearance). In other words, the numerator (denominator) quantifies to what extent vaccine-tvpe infection accelerates or decelerates acquisition (clearance) of non-vaccine-type infection. As a whole, the ratio captures the joint effects of interactions both in acquisition and clearance on the occurrence of non-vaccine-type infection.

Of note, the above pairwise odds ratio (1) has been used for inferring interactions between HPV genotypes [7, 25, 26, 27], whereas the pairwise cross hazard ratio (2) has been used for describing competition between *Streptococcus pneumoniae* serotypes [28].

### 2.4 Predictors in a multi-type setting

#### 2.4.1 Type-specific cross hazard ratio as a predictor of type-specific replacement

We now generalize the pairwise cross hazard ratio (2) to a setting with an arbitrary number of interacting vaccine and non-vaccine types. To predict replacement by a given non-vaccine i Table 1 (see Figure 1a).

**Table 1:**
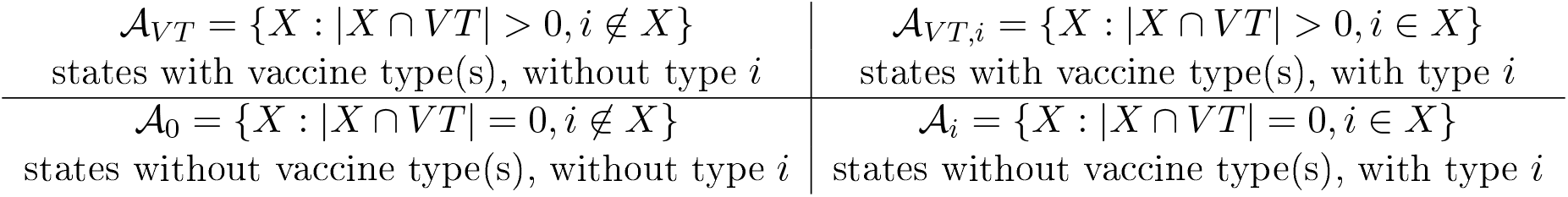
Four disjoint collections of infection states constructed for the definition of the type-specific cross hazardratio (3). Here, *VT* denotes the set of vaccine types.

Based on the transitions between these four aggregate states, we propose the following generalization of the four hazards in the pairwise cross hazard ratio (2):

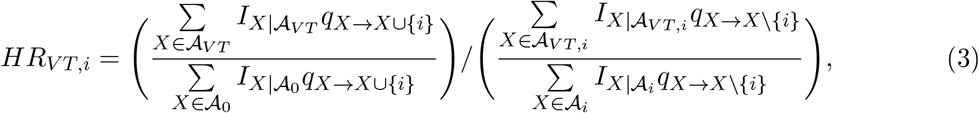

where 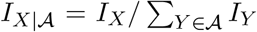 denotes the relative prevalence of state *X* conditioned within aggregate state 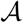. Each term in expression (3) is a weighted average of hazards as it aggregates hazards of acquiring or clearing type i with weights given by the conditional steady-state prevalence of the states from which the transitions occur. In effect, this weighting adjusts for the time each individual spends being at risk for the respective transitions [29]. Similarly to the pairwise cross hazard ratio (2), the numerator and denominator of the type-specific cross hazard ratio (3) are hazard ratios of acquiring and clearing type i in those infected with at least one vaccine type (|*X* ∩ *VT*| > 0) versus those uninfected with any of the vaccine types (|*X* ∩ *VT*| = 0), respectively. In short, the type-specific cross hazard ratio (3) combines interactions by the vaccine types on the non-vaccine type of interest.

##### Example with multiple vaccine types

Assume that *VT* = {1, 2} *NVT* = {3}, and there is interaction only in acquisition (Figure 1a), and consider the type-specific cross hazard ratio (3) as a predictor for replacement by type 3. As the denominator of expression (3) now equals one, the predictor reduces to

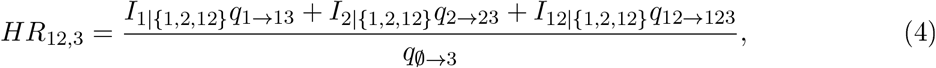

where 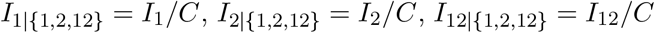 and *C* = *I*_1_ + *I*_2_ + *I*_12_.

#### 2.4.2 Type-specific odds ratio as a predictor of type-specific replacement

The type-specific cross hazard ratio *HR_VT,i_* as given by (3) requires estimation of type-specific acquisition and clearance hazards from longitudinal data. However, collecting such data may be cumbersome and expensive due to repeated observations of the infection states in the same study subjects. It would thus be advantageous if the cross hazard ratio *HR_VT,i_* could be approximated using steady-state cross-sectional (i.e. prevalence) data only, in a similar way as in the two-type setting using the pairwise odds ratio (1).

In a setting with more than two types, the pairwise odds ratio (1) can be generalized to the following type-specific odds ratio:

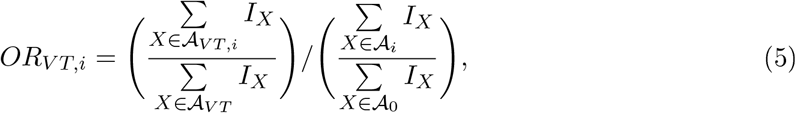

with any of the vaccine types. The approximation of *HR_VT,i_* by *OR_VT,i_* is exact given detailed balance, i.e. when *I_XqX→Y_* = *I_YqY→X_* for any pair of states *X, Y* (see Section A of electronic supplementary material for the proof). In reality, whether this property holds depends on the underlying structure of type interactions.

##### Example continued

In the previous example where *VT* = {1, 2} *NVT* = {3} (Figure 1a), the type-specific odds ratio (5) becomes

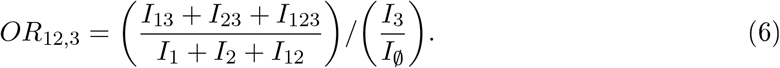

#### 2.4.3 Overall odds and cross hazard ratios as predictors of overall type replacement

Pooling across pairwise odds ratios has been used to summarize interactions of all non-vaccine types with each vaccine type separately [7, 25]. However, this approach lacks a clear interpretation for predicting type replacement. Here, we propose the following overall odds ratio for predicting overall replacement:

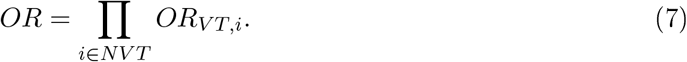

The use of a pooled odds ratio for prediction can be justified by considering the ratio of the odds of overall non-vaccine-type infection in the pre-versus post-vaccination steady states. This odds ratio is essentially the true impact of vaccination with value less than one characterizing overall replacement. Assuming mutual independence between all non-vaccine types, this true odds ratio of overall infection can be approximated by the product of true odds ratios of type-specific infections (see Section B of electronic supplementary material for the derivation). As each true type-specific odds ratio can be predicted by the corresponding *OR_VT,i_*, we envision their product (7) to be a predictor for overall replacement.

Owing to the relation between *HR_VT,i_* and *OR_VT,i_*, we also propose the following overall cross hazard ratio as a predictor for overall replacement:

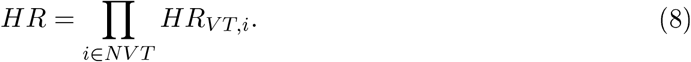

### 2.5 Simulated structures of type interactions

Thus far, we have made no assumptions about the structure of type interactions, i.e. no constraints on *k_Xi_* and *h_Xi_* in defining 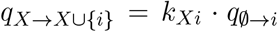 and 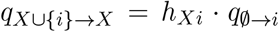. To investigate the performance of the proposed predictors and their robustness against different interaction structures, we considered the following three alternative structures in simulations (see Section C of electronic supplementary material for their exact descriptions).

The first interaction structure is pairwise-symmetric and multiplicative so that each (co) infecting type contributes multiplicatively to the acquisition hazard of an incoming type, i.e. 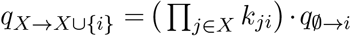 or to the clearance hazard of a coinfecting type, i.e. 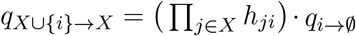. Here *k_ji_* and *h_ji_* describe how (co)infection with type *j* modifies acquisition and clearance of type *i*, respectively. Values of *k_ji_* and 1/*h_ji_* equal to, less, and larger than one correspond to independent, competitive, and synergistic interactions, respectively. In order words, competition decreases the acquisition rate, whereas synergy decreases the clearance rate. Furthermore, pairs of interaction parameters are assumed to be identical, i.e. *k_ij_* = *k_ji_ h_ij_* = *h_ji_*, symmetric interaction matrices. With n types, the numbers of parameters governing the interaction in acquisition and clearance are both reduced from *n*(2^*n*-1^ – 1) to *n*(*n* – 1)/2 (Figure 1b). This pairwise-symmetric multiplicative structure preserves the detailed balance property (see Section D of electronic supplementary material for the proof), so that *HR_VT,i_* md *OR_VT,i_* are equivalent.

Departure from the above structure may disrupt this equivalence of *HR_VT,i_* and *OR_VTyi_.* Two alternative structures are the pairwise-asymmetric multiplicative and the groupwise-symmetric multiplicative structure. The pairwise-asymmetric multiplicative structure relaxes the symmetry constraint *k_ji_* = *k_ij_*. The groupwise-symmetric multiplicative structure assumes the multi-plicativitv to act per groups of types instead of individual types. For example, in a four-type system with groups *A* = {1, 2} *B* = {3, 4}, and interaction parameters *k_A_, k_B_* (within groups), *k_AB_* = *k_BA_* (between groups), the hazards of acquiring type 1 from state 23 and 234 are both 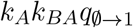.

### 2.6 Performance analysis

To evaluate the performance of the proposed predictors, we simulated the effect of vaccination using a deterministic SIS transmission model (see Section C of electronic supplementary material for details of the model). We investigated how the performance of the predictors depends on the numbers of vaccine and non-vaccine types and their interaction structure. For each setting, different sets of model parameters, including interaction parameters and type-specific transmissibilitv, were randomly generated.

The interaction parameters of the two symmetric structures were uniformly generated on a log scale in the interval (1/3, 3), ranging from competitive to synergistic interactions. To examine the effect of increasing asymmetry under the pairwise-asymmetric structure, asymmetric interaction parameters 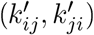 were obtained bv either perturbing the generated parameters of the pairwise-symmetric structure (*k_ij_*), or by generating new 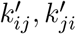 independently of each other. Perturbation of the pairwise-symmetric parameters was done by adding deviations on a log scale, i.e. 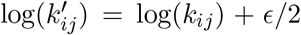 and 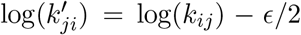. In effect, *e^ϵ^* is the ratio between a pair of reverse interaction parameters, 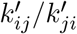 with increasing *ϵ* inducing more asymmetry. When generating 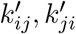 independently, there was no constraint on their ratio.

For each parameter set, prediction of replacement was made at the simulated pre-vaccination steady state and compared to the ‘true’ outcome (replacement/no replacement) at the postvaccination steady state. For each setting, the performance of each predictor was defined as the proportion of correct predictions among all generated parameter sets (see Section E of electronic supplementary material for the exact simulation procedure).

## 3 Simulation results

### 3.1 Performance under the pairwise-symmetric multiplicative structure

As the pairwise-symmetric multiplicative structure obeys detailed balance, the hazard-based and odds-based predictors are equivalent (*HR_VT,i_* = *OR_VT,i_* and *HR* = *OR*) and indeed performed identically (row 1 of Figure 2).

**Figure 2:**
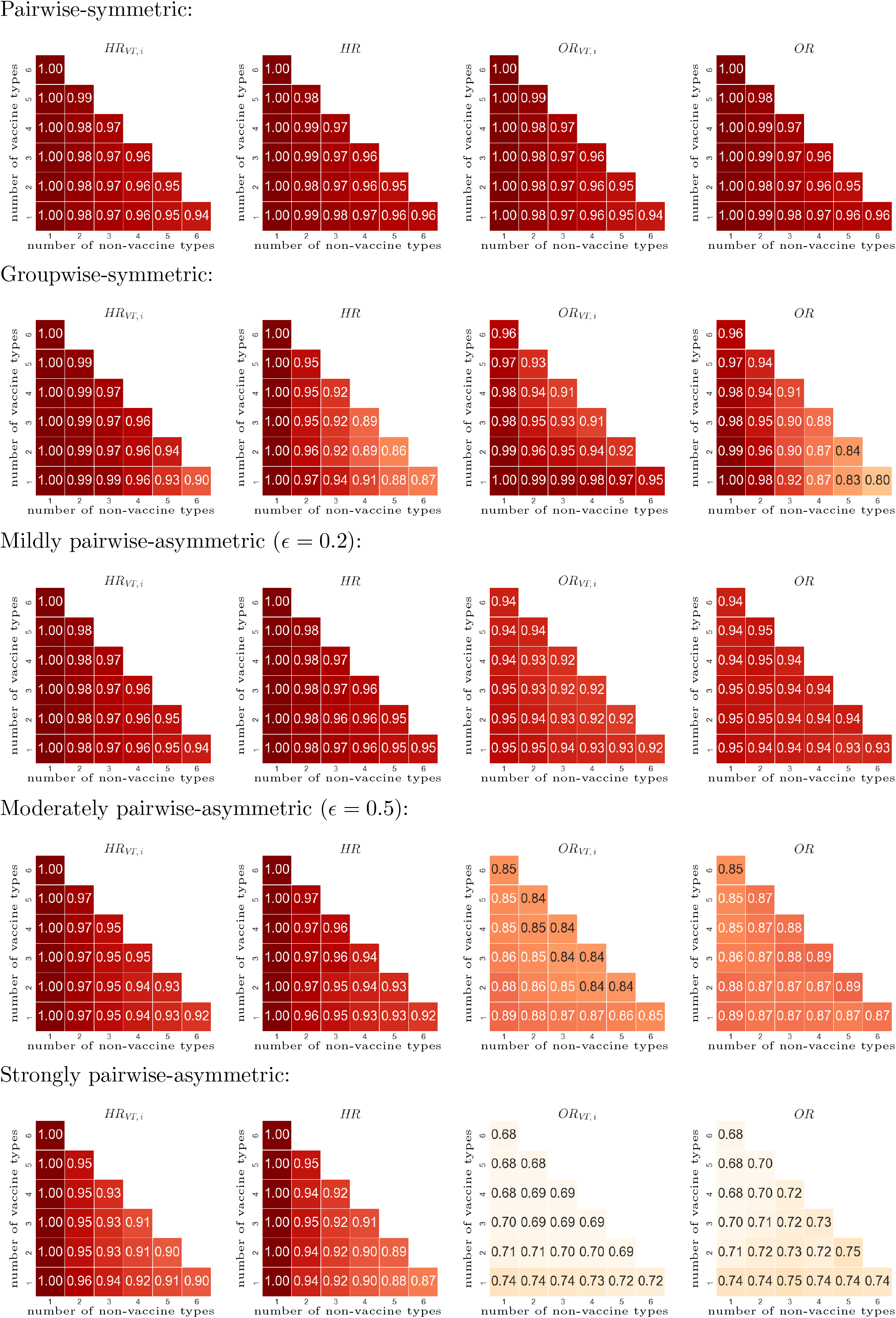
Performance (proportion of correct predictions among all generated parameter sets) of predictors *HR_VT,i_*, HR, *OR_VT,i_*, and *OR* under alternative multiplicative structures for interaction in acquisition (and no interaction in clearance). Row 1: pairwise-symmetric. Row 2: groupwise-symmetric. Rows 3-5: increasing pairwise-asymmetric with increasing values of e for more asymmetry between pairs of reciprocal interaction parameters. Performance of *HR_VT,i_* and *OR_VT,i_* obtained by averaging over the performance of each non-vaccine type *i*.

With one non-vaccine type, the predictors (*HR_VT,i_* = *OR_VT,i_* = *HR* = *OR* correct, even when some vaccine types would interact synergistically while others would compete with the non-vaccine type. Thus the predictors correctly captured the balance between the opposing forces of competition and synergy.

With multiple non-vaccine types, prediction of both type-specific and overall replacement became more difficult due to interactions between the non-vaccine types. For example, in a system with *VT* = {1} *NVT* = {2, 3}, if *k*_13_ = 1, *k*_12_ > 1 and *k*_23_ < 1 (Figure 1b), vaccination indirectly triggered replacement by type 3 by decreasing type-2 infection and thus increasing type-3 infection. Rewriting expression (3) shows that this indirect effect is indicated by the second factor in

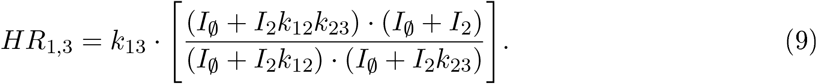

In particular, the expression in the brackets is less than one when either *k*_12_ or *k*_23_ (but not both) is less than one (see Section F of electronic supplementary material for the derivation of (9)). However, if the direct and indirect effects acted in opposite directions, the predictors did not always adequately capture their combined effect (see Section G of electronic supplementary material for an example). Indeed, with multiple non-vaccine types, the performance decreased as the number of non-vaccine types increased (follow each subfigure in Figure 2 across the ↘ diagonals). Nonetheless, the performance of all predictors remained well above 94%. In addition, when the overall predictor failed with a value close to one, the overall prevalence of non-vaccine-tvpe infection usually changed only modestly (see Section G of electronic supplementary material for an example).

### 3.2 Performance under alternative interaction structures

Under the alternative interaction structures, the hazard-based and odds-based predictors were no longer equivalent but still performed comparably, except when the asymmetry was strong (Figure 2). With one non-vaccine type, the hazard-based predictions remained almost perfect unlike the odds-based ones. Under the groupwise-symmetric structure, the predictive performance decreased more rapidly as the number of non-vaccine types increased, as compared to the pairwise-symmetric structure. However, it was still above 80% in simulations with up to seven types (row 2 of Figure 2).

Under the pairwise-asymmetric structure, the performance of both sets of predictors decreased with increasing asymmetry (row 3 to 5 of Figure 2). Nevertheless, the performance of the hazard-based predictors remained above 85% even when pairs of *k_ij_* and *k_ji_* were generated independently, inducing strong asymmetry. The decreasing performance was more vivid for the odds-based predictors, which went down to 70% in the corresponding settings. The superiority of the hazard-based predictors can be illustrated in the following example of two types. Suppose that the vaccine type competes with the non-vaccine type (*k*_12_ < 1) while the reverse interaction is synergistic (*k*_21_ > 1). Then *HR*_1,2_ = *k*_12_ < 1 predicts replacement correctly, whereas the symmetrically defined *OR*_1,2_ may not because it averages over *k*_12_ and *k*_21_.

The above results were based on simulations in which interactions occurred only in acquisition. Additional simulations that allowed interactions in both acquisition and clearance showed almost identical performance of both the hazard-based and odds-based predictors (compare Figure 2 to Section H of electronic supplementary material).

## 4 Discussion

This paper aims to develop methodology for predicting type replacement for pathogens with many, potentially interacting types using pre-vaccination epidemiological data. We have proposed a predictor of replacement by individual non-vaccine types (i.e. increase in type-specific prevalence) and another predictor of increase in the overall non-vaccine-type prevalence. Both predictors are initially defined in terms of hazards of acquiring and clearing the non-vaccine types and thus, in principle, require estimation of longitudinal data. In addition, we have derived alternative predictors that can be readily estimated from cross-sectional prevalence data as odds ratios. All proposed predictors demonstrated satisfactory performance (mostly above 85%) across a range of simulated structures of type interactions except when the interactions were strongly asymmetric.

The predictor of replacement by a given non-vaccine type is constructed as a weighted cross hazard ratio of acquiring versus clearing infection with the non-vaccine type in question, comparing those infected and uninfected with any of the vaccine types. The odds-based alternative is defined as the odds of infection with that non-vaccine type in presence versus absence of infection with any of the vaccine types.

We note the similarity between type interaction and vaccine efficacy of multi-valent vaccines as both are defined in terms of cross ratios of weighted hazards in a multi-type setting. In both cases, it proves useful to weigh hazards with the steady-state prevalence, conditioned on appropriate risk sets. In defining estimators for vaccine efficacy against one or more vaccine types, the risk sets are those from which acquisitions of the vaccine types may occur [29]. Likewise, in the current paper, the risk sets are those from which acquisitions (or clearances) of the non-vaccine type may occur (see the type-specific cross hazard ratio (3)).

The predictors of overall replacement take form as products of either the hazard-based or the odds-based type-specific predictors. In effect, the overall predictors are obtained by pooling the type-specific predictors on a logarithmic scale. In various studies that have used pairwise odds ratios to evaluate HPV type interactions, pooling has been performed across the pool of non-vaccine types for each vaccine type separately [7, 25, 26]. These pooled odds ratios have been interpreted as the affinity of a given vaccine type to be involved in coinfection with any of the non-vaccine types, but their relevance for predicting type replacement has remained elusive. Our results substantiate the predictive value of pooled odds ratios regarding overall replacement, but suggest that pooling is better performed on the entire set of vaccine types, instead of for each vaccine type separately. Furthermore, the pairwise odds ratios have been compared to the pooled ones to identify likely candidates for type replacement [7, 25, 26]. Accordingly, the potential for replacement by a given non-vaccine type has been assessed separately for each vaccine type, while our type-specific predictors capture interactions with all vaccine types in a comprehensive way.

Our simulation study revealed how the applicability of the new predictors depends on the underlying structure of type interactions. The predictors performed best under the pairwise-symmetric multiplicative structure, in which the hazard-based and odds-based predictors are equivalent. The predictors were mostly able to capture the opposing forces of competition and synergy as well as the interplay between interactions in acquisition and clearance. Under other simulated structures, the hazard-based predictors still performed well, while the odds-based predictors performed fairly up to a reasonable degree of asymmetry. As a rule of thumb, symmetric interactions facilitate prediction of type replacement, while complex and heterogeneous interactions may necessitate more sophisticated predictors that capture details and directions of interactions. For HPV, little is known about the structure of interactions between genotypes as the very existence of interaction is already difficult to determine. For instance, any imbalance between type-specific prevalence may mask possible asymmetric interactions, and although pooling of multiple types may increase the power of detecting interaction, it may also obscure type-specific patterns. Nevertheless, our simulation study demonstrated that the new predictors are robust against various interaction structures.

We cannot ensure good performance of the proposed predictors under mechanisms of interactions other than the ones we considered. For instance, if the pathogen types interact through natural cross-immunity that is long-lasting, infections with different types may be positively associated (i.e. as given by odds ratios greater than one) while there is a risk for type replacement [30, 31]. This shortcoming is inherent to the difficulty to distinguish between susceptible individuals and those who have acquired natural immunity, as is the case for HPV infection which induces only a weak antibody response [32]. Nevertheless, if immunity is confined to be type-specific, the predictors work equally well in a two-type model [21], and we envision the proposed predictors to remain applicable also in a multi-type setting. Another mechanism that we did not consider is competition for transmissibility, e.g. through reduction of viral load during coinfection [13, 14, 15]. Although competition for transmissibility is not captured by the proposed predictors, it is likely correlated with competition in clearance, as HPV persistence is also determined by viral load [16]. Hence, the predictors may retain good predictive ability even when not all mechanisms of interactions are captured.

Despite the theoretical appeal of the proposed predictors and their promising performance in simulations, some challenges still need to be addressed. First, more extensive simulations are needed to investigate how well the predictors perform with high numbers of pathogen types. Here, simulations were performed with at most seven types, while HPV consists of up to fifteen high-risk types. Second, our method assumes elimination of all vaccine-targeted types, although some vaccine types or cross-protective types may persist, especially when vaccination coverage is low [4, 5]. If persistence of targeted types would mostly prevent or limit the extent of type replacement, our method may still provide good predictions as a worst-case scenario. However, formal analysis would be required to sharpen this intuition. In addition, we ignore evolution of the model pathogen types for the timescale of type replacement, which seems plausible for HPV given its low mutation rate [33]. However, in the long run, vaccination may induce an evolutionary change of the vaccine types, which could lead to type replacement by novel (as opposed to evolutionarily stable) types. Third, it remains to be investigated whether the hazard-based and odds-based predictors can be accurately estimated from limited data. In particular, although the hazard-based predictors performed better in simulations, it is not straightforward which ones are more suitable for the empirical setting. In general, hazard-based estimates are more robust against different sources of bias (e.g. due to population entry and unobserved heterogeneity) [31] but require a larger sample size than estimation of type-specific point prevalence. In addition, statistical methods need to be developed to deal with possible confounding due to common risk factors, which were neglected in our analysis. In particular, multivariate statistical methods (e.g. GEE and random effect models) may be adapted to adjust for observed heterogeneity or control for unobserved heterogeneity when estimating the new predictors, which are essentially odds and hazard ratios.

For HPV, most epidemiological studies have concluded a low risk of type replacement based on a lack of systematic patterns of negative associations in the co-occurrence of vaccine and non-vaccine types [12, 25]. The methodology presented in this paper may help to translate the observed patterns into explicit prediction of the overall risk of replacement. Additionally, application of our type-specific predictors could discover hitherto hidden potential for type replacement, as this potential is not only determined by direct interactions with the vaccine types but also shaped though indirect interactions with other non-vaccine types. As the circulation of vaccine types will be further reduced in the near future, more data will become available to validate any predictions about the long-term impact of HPV vaccination. The proposed methodology could also be used to give better insights into the underpinnings of type competition in *Streptococcus pneumoniae*, a pathogen for which replacement has been widely observed after the introduction of the pneumococcal conjugate vaccination [34, 35].

To conclude, we developed novel methodology for predicting type replacement in a setting of many interacting types. We did so by relating available epidemiological data to the underlying mechanisms of how pathogen types may interact. The proposed predictors may help to better anticipate and understand the impact of vaccination against pathogens with many coexisting (sub)tvpes.

## Supporting information

Electronic Supplementary Material

