## Supplementary material for "Capturing multiple-type interactions into practical predictors of type replacement following HPV vaccination": Electronic Supplementary Material

### A Equivalence of hazard-based and odds-based predictors under detailed balance

We prove the equivalence of the type-specific hazard-based and odds-based predictors (defined in (3) and (5) in the main text, respectively) under detailed balance, i.e. assuming that $I_X q_{X\to Y} = I_Y q_{Y\to X}$ for any pair of states $X, Y$. According to definition (3),

$$HR_{VT,i} = \left( \frac{\sum\limits_{X\in\mathcal{A}_{VT}} I_{X|\mathcal{A}_{VT}} q_{X\to X\cup\{i\}}}{\sum\limits_{X\in\mathcal{A}_0} I_{X|\mathcal{A}_0} q_{X\to X\cup\{i\}}} \right) / \left( \frac{\sum\limits_{X\in\mathcal{A}_{VT,i}} I_{X|\mathcal{A}_{VT,i}} q_{X\to X\setminus\{i\}}}{\sum\limits_{X\in\mathcal{A}_i} I_{X|\mathcal{A}_i} q_{X\to X\setminus\{i\}}} \right)$$

$$= \left( \frac{\frac{\sum_{X\in\mathcal{A}_{VT}} I_X q_{X\to X\cup\{i\}}}{\sum_{X\in\mathcal{A}_{VT}} I_X}}{\frac{\sum_{X\in\mathcal{A}_0} I_X q_{X\to X\cup\{i\}}}{\sum_{X\in\mathcal{A}_0} I_X}} \right) / \left( \frac{\frac{\sum_{X\in\mathcal{A}_{VT,i}} I_X q_{X\to X\setminus\{i\}}}{\sum_{X\in\mathcal{A}_{VT,i}} I_X}}{\frac{\sum_{X\in\mathcal{A}_i} I_X q_{X\to X\setminus\{i\}}}{\sum_{X\in\mathcal{A}_i} I_X}} \right),$$

where each of the four aggregate-level hazards are specified as wegihted averages of the respective incoming type-specific hazards (see text). Reordering the terms as $\left(\frac{\frac{a_1}{a_2}}{\frac{b_1}{b_2}}\right) / \left(\frac{\frac{c_1}{c_2}}{\frac{d_1}{d_2}}\right) = \left(\frac{a_1\cdot b_2}{a_2\cdot b_1}\right)\cdot\left(\frac{c_2\cdot d_1}{c_1\cdot d_2}\right) = \left(\frac{a_1\cdot d_1}{b_1\cdot c_1}\right)\cdot\left(\frac{b_2\cdot c_2}{a_2\cdot d_2}\right)$ gives

$$HR_{VT,i} = \left( \frac{\sum\limits_{X\in\mathcal{A}_{VT}} I_X q_{X\to X\cup\{i\}} \cdot \sum\limits_{X\in\mathcal{A}_i} I_X q_{X\to X\setminus\{i\}}}{\sum\limits_{X\in\mathcal{A}_0} I_X q_{X\to X\cup\{i\}} \cdot \sum\limits_{X\in\mathcal{A}_{VT,i}} I_X q_{X\to X\setminus\{i\}}} \right) \cdot \left( \frac{\sum\limits_{X\in\mathcal{A}_0} I_X \cdot \sum\limits_{X\in\mathcal{A}_{VT,i}} I_X}{\sum\limits_{X\in\mathcal{A}_{VT}} I_X \cdot \sum\limits_{X\in\mathcal{A}_i} I_X} \right).$$

As each state in $\mathcal{A}_i$ can be reached from $\mathcal{A}_0$ by adding (acquiring) type $i$, the sum over $\mathcal{A}_i$ can be written as a sum over $\mathcal{A}_0$ (and analogously for $\mathcal{A}_{VT,i}$ and $\mathcal{A}_{VT}$):

$$HR_{VT,i} = \left( \frac{\sum\limits_{X\in\mathcal{A}_{VT}} I_X q_{X\to X\cup\{i\}} \cdot \sum\limits_{X\in\mathcal{A}_0} I_{X\cup\{i\}} q_{X\cup\{i\}\to X}}{\sum\limits_{X\in\mathcal{A}_0} I_X q_{X\to X\cup\{i\}} \cdot \sum\limits_{X\in\mathcal{A}_{VT}} I_{X\cup\{i\}} q_{X\cup\{i\}\to X}} \right) \cdot \left( \frac{\sum\limits_{X\in\mathcal{A}_0} I_X \cdot \sum\limits_{X\in\mathcal{A}_{VT,i}} I_X}{\sum\limits_{X\in\mathcal{A}_{VT}} I_X \cdot \sum\limits_{X\in\mathcal{A}_i} I_X} \right).$$

Finally, invoking detailed balance shows that $HR_{VT,i}$ equals $OR_{VT,i}$:

$$HR_{VT,i} = \left( \frac{\sum\limits_{X\in\mathcal{A}_{VT}} I_X q_{X\to X\cup\{i\}} \cdot \sum\limits_{X\in\mathcal{A}_0} I_X q_{X\to X\cup\{i\}}}{\sum\limits_{X\in\mathcal{A}_0} I_X q_{X\to X\cup\{i\}} \cdot \sum\limits_{X\in\mathcal{A}_{VT}} I_X q_{X\to X\cup\{i\}}} \right) \cdot \left( \frac{\sum\limits_{X\in\mathcal{A}_0} I_X \cdot \sum\limits_{X\in\mathcal{A}_{VT,i}} I_X}{\sum\limits_{X\in\mathcal{A}_{VT}} I_X \cdot \sum\limits_{X\in\mathcal{A}_i} I_X} \right)$$

$$= \left( \frac{\sum\limits_{X\in\mathcal{A}_{VT,i}} I_X}{\sum\limits_{X\in\mathcal{A}_{VT}} I_X} \right) / \left( \frac{\sum\limits_{X\in\mathcal{A}_i} I_X}{\sum\limits_{X\in\mathcal{A}_0} I_X} \right) = OR_{VT,i}.$$

#### B Pooled predictor of overall replacement

We here prove that the odds ratio of overall non-vaccine-type infection ($\overline{OR}_{NVT}$), comparing the pre- and post-vaccination steady states, can be approximated by the product of the odds ratios of non-vaccine-type-specific infections, $\prod_{i \in NVT} \overline{OR}_i$, under mutually independent non-vaccine-type infection dynamics. We start by introducing some notations:

- $I_X$: pre-vaccination steady-state prevalence of infection state $X$;
- $I'_X$: post-vaccination steady-state prevalence of infection state $X$;
- $E_i = \sum_{i \in X} I_X$: pre-vaccination prevalence of type-$i$ infection;
- $E'_i = \sum_{i \in X} I'_X$: post-vaccination prevalence of type-$i$ infection;
- $odds_i = E_i/(1 - E_i)$: pre-vaccination odds of type-$i$ infection;
- $odds'_i = E'_i/(1 - E'_i)$: post-vaccination odds of type-$i$ infection;
- $\overline{OR}_i = odds_i/odds'_i$: true odds ratio of type-$i$ infection, comparing the pre- and post-vaccination steady states;
- $E_{NVT} = \sum_{|X \cap NVT|>0} I_X$: pre-vaccination overall non-vaccine-type prevalence;
- $E'_{NVT} = \sum_{|X \cap NVT|>0} I'_X$: post-vaccination overall non-vaccine-type prevalence;
- $odds_{NVT} = E_{NVT}/(1 - E_{NVT})$: pre-vaccination odds of overall non-vaccine-type infection;
- $odds'_{NVT} = E'_{NVT}/(1 - E'_{NVT})$: post-vaccination odds of overall non-vaccine-type infection;
- $\overline{OR}_{NVT} = odds_{NVT}/odds'_{NVT}$: true odds ratio of overall non-vaccine-type infection, comparing the pre- and post-vaccination steady states;

For notational convenience, we have purposefully defined the above odds ratios with the pre-vaccination odds in the numerator and the post-vaccination odds in the denominator. Accordingly, $\overline{OR}_i < 1$ and $\overline{OR}_{NVT} < 1$ characterize type-specific and overall replacement, respectively, corresponding to the way values less than one of $OR_i$ and $OR$ predict replacement.

We now prove that $\overline{OR}_{NVT} \approx \prod_{i \in NVT} \overline{OR}_i$ for two non-vaccine types. The proof for more than two non-vaccine types proceeds analogously. We start by rewriting $odds_{NVT}$:

$$odds_{NVT} = \frac{E_{NVT}}{1 - E_{NVT}} = \frac{I_1 + I_2 + I_{12}}{I_\emptyset}.$$

By assuming independence between the non-vaccine types, i.e. $I_{12} = E_1E_2$:

$$\begin{aligned} odds_{NVT} &= \frac{E_1(1-E_2) + (1-E_1)E_2 + E_1E_2}{(1-E_1)(1-E_2)} \\ &= \frac{E_1}{1-E_1} + \frac{E_2}{1-E_2} + \frac{E_1E_2}{(1-E_1)(1-E_2)} \\ &= odds_1 + odds_2 + odds_1 odds_2 \\ &= \overline{OR}_1 odds'_1 + \overline{OR}_2 odds'_2 + \overline{OR}_1\overline{OR}_2 odds'_1 odds'_2. \end{aligned}$$

Similarly, $odds'_{NVT} = odds'_1 + odds'_2 + odds'_1 odds'_2$. The true odds ratio for the overall non-vaccine-type infection then is

$$\overline{OR}_{NVT} = odds_{NVT}/odds'_{NVT} = \frac{\overline{OR}_1 odds'_1 + \overline{OR}_2 odds'_2 + \overline{OR}_1\overline{OR}_2 odds'_1 odds'_2}{odds'_1 + odds'_2 + odds'_1 odds'_2},$$

which has a value less than one when overall type replacement occurs. As the term $\overline{OR}_1\overline{OR}_2$ appears in the expression of the true odds ratio, $\overline{OR}_{NVT}$, it can be used as a predictor of overall type replacement. How well $\overline{OR}_{NVT}$ and $\overline{OR}_1\overline{OR}_2$ correspond to one another can be illustrated in the following example. eFigure 1 shows for which combination of values of $\overline{OR}_1$ and $\overline{OR}_2$ the predictor, $\overline{OR}_1\overline{OR}_2$, is less than one (the area below the dashed-dotted curve), predicting replacement. The area in which the true odds ratio of overall non-vaccine-type prevalence is less than one (the area below the dashed curve) is seen to coincide, except for small areas indicated by the arrows in the figure. The figure was generated with $odds_1 = 0.64$ and $odds_2 = 0.37$. Other values give similar results. Due to this correspondence, $\overline{OR}_{NVT}$ can be well approximated by $\overline{OR}_1\overline{OR}_2$.

As, in addition, each $\overline{OR}_i$ can be predicted by $OR_{VT,i}$ (defined in (5) in the main text), we envision that $\overline{OR}_{NVT}$ can be well predicted by $\prod_{i \in NVT} OR_{VT,i}$ (defined in (7) in the main text). The performance of using $\prod_{i \in NVT} OR_{VT,i}$ to predict overall replacement is evaluated in Section 3 of the main text.

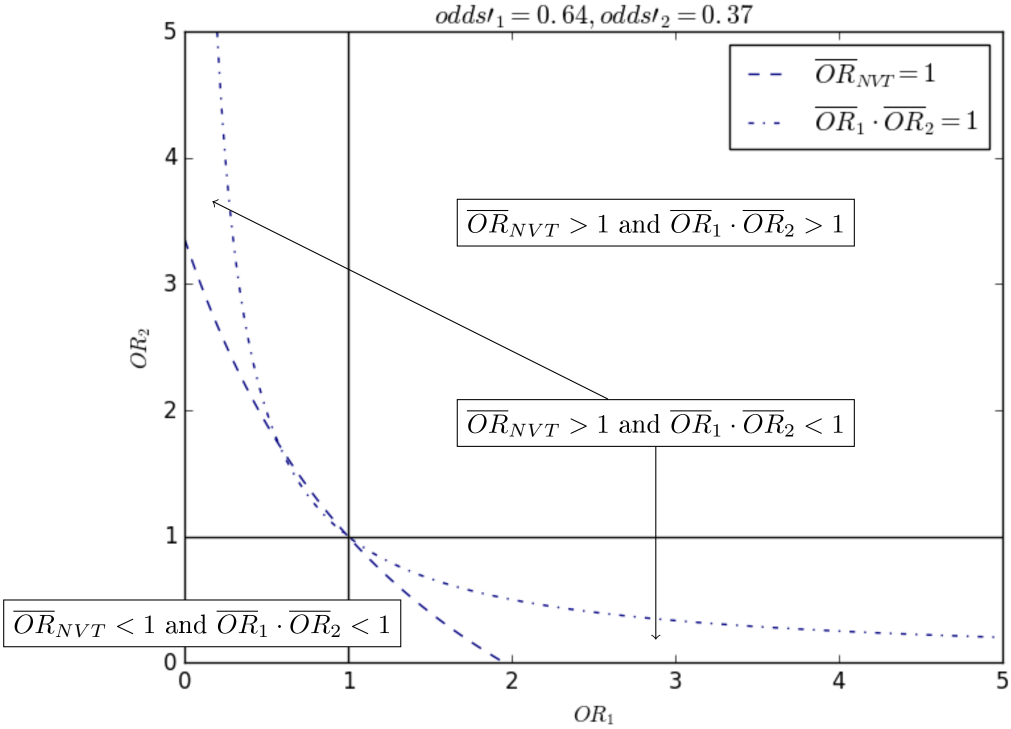


eFigure 1: Contour plot of $\overline{OR}_{NVT}$ and $\overline{OR}_1\overline{OR}_2$ with contour lines at $\overline{OR}_{NVT} = 1$ and $\overline{OR}_1\overline{OR}_2 = 1$, respectively. The values of $odd'_1 = 0.64$ and $odd'_2 = 0.37$ were randomly generated.

### C A multi-type Susceptible-Infected-Susceptible model

In this section, we describe the transmission model used to simulate the impact of vaccination. The model has Susceptible-Infected-Susceptible dynamics with regard to each of the pathogen types. It allows interactions in both acquisition and clearance. We describe the model's infection states and the respective transition hazards in Section C.1, the pre- and post-vaccination steady states in Section C.2, and three alternative structures of type interactions in Section C.3.

#### C.1 Infection states and transition hazards

For a pathogen with $n$ types, the model consists of $2^n$ infection states, each defined by the set of types the host is infected with. We denote the set of infection states by $\mathcal{S}$. Acquisitions and clearances of type-specific infection are assumed to be sequential so that only transitions between states that differ by infection of one type are possible. Figure1a in the main text shows the eight infection states and the respective transitions for a pathogen with three types.

The baseline hazard for a susceptible individual to acquire type $i$ is $q_{\emptyset\to i} = c\beta_i \sum_{i\in X} I_X$, where $c$ is the contact rate, and $\beta_i$ is the proportion (probability) of successfully acquiring type $i$ given contact with an individual infected with type $i$. The baseline hazard of clearing type $i$ by a singly infected individual is $q_{i\to\emptyset}$. The other hazards of acquisition and clearance are given by

$$\begin{cases} q_{X\to X\cup\{i\}} = k_{Xi} \cdot q_{\emptyset\to i}, \\ q_{X\cup\{i\}\to X} = h_{Xi} \cdot q_{i\to\emptyset}, \end{cases}$$

where $k_{Xi}$ and $h_{Xi}$ are defined by the underlying structure of type interaction (cf. Section C.3).

#### C.2 System of ordinary differential equations

A system of ordinary differential equations describes how the proportions of individuals in each of the model states evolve over time. The equation for state $X$ is

$$\frac{dI_X}{dt} = \sum_{i\in X} I_{X\setminus\{i\}} q_{X\setminus\{i\}\to X} - \sum_{i\in X} I_X q_{X\to X\setminus\{i\}} + \sum_{i\notin X} I_{X\cup\{i\}} q_{X\cup\{i\}\to X} - \sum_{i\notin X} I_X q_{X\to X\cup\{i\}},$$

where the four terms correspond to:

- transition from $I_{X\setminus\{i\}}$ to $I_X$ by acquisition of type $i \in X$;
- transition from $I_X$ to $I_{X\setminus\{i\}}$ by clearance of type $i \in X$;
- transition from $I_{X\cup\{i\}}$ to $I_X$ by clearance of type $i \notin X$;

- transition from $I_X$ to $I_{X\cup\{i\}}$ by acquisition of type $i \notin X$.

The pre-vaccination steady state $\{I_X\}_{X\in\mathcal{S}}$ is defined by the condition $\frac{dI_X}{dt} = 0$. The post-vaccination steady state $\{I'_X\}_{X\in\mathcal{S}}$ is determined by the condition $\frac{dI'_X}{dt} = 0$ and the condition $I'_X = 0$ for any state containing vaccine types, i.e. $|X \cap VT| > 0$. In effect, the post-vaccination steady state is defined as the steady state after the elimination of all vaccine types. Thereby, we implicitly assume a vaccination scheme (i.e. coverage of vaccination and efficacy against the vaccine types) that leads to elimination. Note that, in the model, the non-vaccine-type prevalence in such a post-vaccination steady state is the same regardless of the exact vaccination scheme. As long as the scheme achieves elimination, the system will converge to the unique non-vaccine-type-only steady state.

#### C.3 Multiplicative structures of type interaction

##### C.3.1 Pairwise-symmetric and pairwise-asymmetric structures

Under both the pairwise-symmetric and pairwise-asymmetric multiplicative interaction structure (eFigure 2), the hazards of acquisition and clearance in presence of infection with other types are

$$\begin{cases} q_{X\to X\cup\{i\}} = \left(\prod_{j\in X} k_{ji}\right) \cdot q_{\emptyset\to i}, \\ q_{X\cup\{i\}\to X} = \left(\prod_{j\in X} h_{ji}\right) \cdot q_{i\to\emptyset}. \end{cases}$$

In other words, each (co)infecting type contributes a multiplicative factor to the hazard of acquiring (clearing) the incoming (outgoing) type. Moreover, under the symmetry assumption, $k_{ij} = k_{ji}$ and $h_{ij} = h_{ji}$ for all $i \neq j$.

##### C.3.2 Groupwise-symmetric structure

The groupwise-symmetric multiplicative structure (eFigure 3) assumes that the multiplicativity acts per groups of types instead of individual types. Given a system with $n$ types and $m \leq n$ groups of types, $A_1, A_2, \ldots, A_m$. Suppose that type $i$ belongs to group $A_l$. The hazards of acquiring and clearing type $i$ then are

$$\begin{cases} q_{X\to X\cup\{i\}} = \left(\prod_{j=1}^{m} \mathbb{1}\{|X \cap A_j| > 0\} \cdot k_{jl}\right) \cdot q_{\emptyset\to i}, \\ q_{X\cup\{i\}\to X} = \left(\prod_{j=1}^{m} \mathbb{1}\{|X \cap A_j| > 0\} \cdot h_{jl}\right) \cdot q_{i\to\emptyset}. \end{cases}$$

The symmetry assumption now reads $k_{jl} = k_{lj}$ and $h_{jl} = h_{lj}$ for all between-group interaction parameters. Furthermore, we abbreviate all within-group interaction parameters $k_{jj}$ and $h_{jj}$ by $k_j$ and $h_j$, respectively.

Here in the electronic supplementary material, we index groups with numbers, whereas alphabets are used in the main text. The example of the main text reads as follows with the numerical notation:

in a four-type system with groups $A_1 = \{1,2\}$, $A_2 = \{3,4\}$, and interaction parameters $k_1$, $k_2$ (within groups), $k_{12} = k_{21}$ (between groups), the acquisition hazards of type 1 from state 23 and 234 are both $k_1 k_{21} q_{\emptyset\to 1}$.

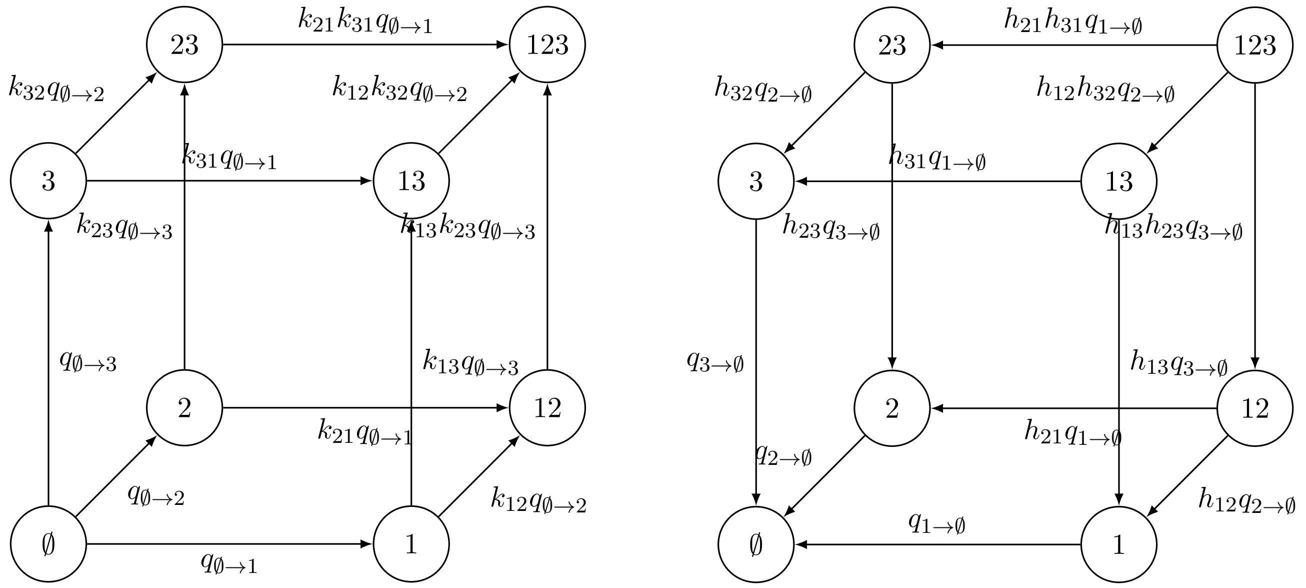


(a) acquisition (b) clearance

eFigure 2: The eight infection states and the respective (a) acquisitions and (b) clearances in a three-type system, under the pairwise-symmetric or pairwise-asymmetric multiplicative structure.

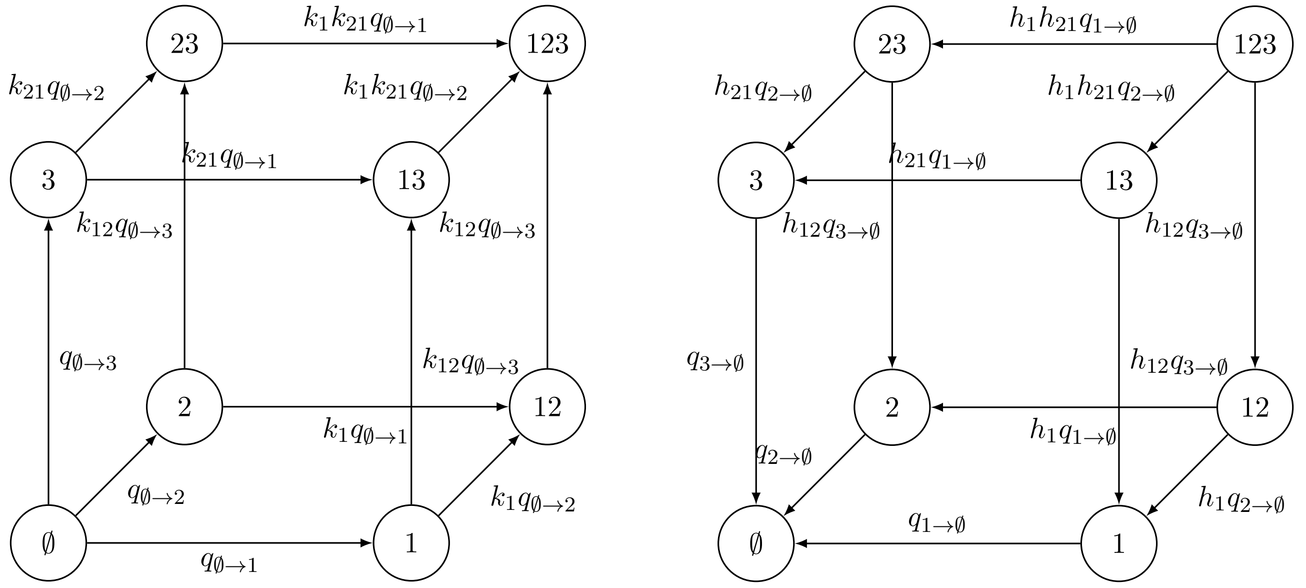


(a) acquisition (b) clearance

eFigure 3: The eight infection states and the respective (a) acquisitions and (b) clearances in a three-type system, under the groupwise-symmetric multiplicative structure with groups $A_1 = \{1,2\}, A_2 = \{3\}$. Here, we index groups with numbers, whereas alphabets are used in the main text.

#### D Detailed balance under the pairwise-symmetric multiplicative structure

We here prove the pairwise-symmetric multiplicative structure as defined in Section C satisfies the detailed balance property, i.e. $I_X q_{X\to Y} = I_Y q_{Y\to X}$ for any pair of states $X, Y$ in the state space $\mathcal{S}$. The proof is based on first proposing a possible steady-state distribution over $\{X\}_{X\in S}$ and then showing that detailed balance holds for this distribution. As detailed balance implies global balance, the distribution is the unique steady-state distribution [1].

We propose the following expression for the steady-state distribution:

$$I_X = \left(\prod_{i\in X} q_{\emptyset\to i}\right)\cdot\left(\prod_{i\notin X} q_{i\to\emptyset}\right)\cdot\left(\frac{\prod_{i<j\in X} k_{ij}}{\prod_{i<j\in X} h_{ij}}\right)/C,\quad X\in\mathcal{S}.$$

Here, $k_{ij}$ and $h_{ij}$ are the interaction parameters, and $C$ is a normalizing constant. Take any pair of states that differ with regard to the status of one type exactly, i.e. $X$ and $Y = X\cup\{e\}$:

$$\begin{aligned}
\frac{I_Y}{I_X} &= \frac{\left(\prod_{i\in Y} q_{\emptyset\to i}\right)\cdot\left(\prod_{i\notin Y} q_{i\to\emptyset}\right)\cdot\left(\frac{\prod_{i<j\in Y} k_{ij}}{\prod_{i<j\in Y} h_{ij}}\right)/C}{\left(\prod_{i\in X} q_{\emptyset\to i}\right)\cdot\left(\prod_{i\notin X} q_{i\to\emptyset}\right)\cdot\left(\frac{\prod_{i<j\in X} k_{ij}}{\prod_{i<j\in X} h_{ij}}\right)/C}\\
&= \frac{q_{\emptyset\to e}\cdot\left(\prod_{i<j\in Y} k_{ij}\right)\cdot\left(\prod_{i<j\in X} h_{ij}\right)}{q_{e\to\emptyset}\cdot\left(\prod_{i<j\in X} k_{ij}\right)\cdot\left(\prod_{i<j\in Y} h_{ij}\right)}\\
&= \frac{q_{\emptyset\to e}\cdot\left(\prod_{i<j\in X} k_{ij}\right)\cdot\left(\prod_{i<j\in X} h_{ij}\right)\cdot\left(\prod_{i\in X} k_{ie}\right)}{q_{e\to\emptyset}\cdot\left(\prod_{i<j\in X} k_{ij}\right)\cdot\left(\prod_{i<j\in X} h_{ij}\right)\cdot\left(\prod_{i\in X} h_{ie}\right)}\\
&= \frac{q_{\emptyset\to e}\cdot\left(\prod_{i\in X} k_{ie}\right)}{q_{e\to\emptyset}\cdot\left(\prod_{i\in X} h_{ie}\right)}\\
&= \frac{q_{X\to Y}}{q_{Y\to X}}.
\end{aligned}$$

For any pair of states that are not connected, $q_{X\to Y} = q_{Y\to X} = 0$ so that $I_X q_{X\to Y} = I_Y q_{Y\to X}$ holds trivially. Hence, we have proved that the proposed distribution satisfies the detailed balance property.

### E Simulation procedure

For any given simulation setting (e.g. the pairwise-symmetric multiplicative structure with $|VT| = 1$ and $|NVT| = 2$), $N = 10000$ different parameter sets were randomly generated. The varying model parameters were the interaction parameters (see below) and the type-specific transmissibilities ($\beta_i$ as defined in Section C.1), while the baseline hazards of clearance were fixed, i.e. $q_{i\to\emptyset} = 1$ for all type $i$. Of note, the results in the main text were generated with no interaction in clearance, i.e. $h_{ij} = 1$. See Section H for the simulation results obtained in the general case with interaction in both acquisition and clearance. The parameter values and ranges used are given in eTable 1.

The varied interaction parameters (as defined in Section C.3) under different interaction structures were generated as follows:

- Pairwise-symmetric multiplicative structure: altogether $n(n-1)/2$ parameters $k_{ij}$ for $i < j$;
- Pairwise-asymmetric multiplicative structure: altogether $n(n-1)$ parameters $k'_{ij}$ for $i \neq j$. These asymmetric interaction parameters were obtained by either perturbing the interaction parameters generated under the pairwise-symmetric multiplicative structure, or by generating $k'_{ij}, k'_{ji}$ independently of each other. Perturbation of the pairwise-symmetric parameters is done by adding or subtracting $\epsilon/2$ on a log scale. Which one of $k'_{ij}$ and $k'_{ji}$ was derived from addition (and which one from subtraction) was decided randomly with 50/50 chance. This construction ensures the order of $k'_{ij}$ and $k'_{ji}$ to be random, while $|\log(k'_{ij}) - \log(k'_{ji})| = \epsilon$ fixed. Hence, the ratio between any pair of reverse interaction parameters $k'_{ij}/k'_{ji}$ (or $k'_{ji}/k'_{ij}$) was $e^{\epsilon} = 1.22, 1.65$. The applied values of $\epsilon$ were 0.2 and 0.5. We also considered a maximal amount of asymmetry by generating $k'_{ij}, k'_{ji}$ independently of each other;
- Groupwise-symmetric multiplicative structure: the number of interaction parameters depended on the randomly generated number of groups $m \leq n$. With $m$ groups, the numbers of parameters were $m$ (for $k_i$) and $m(m-1)/2$ (for $k_{ij}$ where $i < j$).

Interaction parameters under the pairwise-symmetric and the groupwise-symmetric multiplicative structure were generated uniformly on a log scale in the interval $(1/3, 3)$ by means of Latin hypercube sampling, likewise for the pairwise-asymmetric multiplicative structure when generating $k'_{ij}, k'_{ji}$ independently. The type-specific transmissibilities were generated uniformly in the interval $(2.5, 3.5)$ also by means of Latin hypercube sampling.

Note that some generated parameter sets were excluded to retain meaningful simulations. Given a number of types in a setting, we included only parameter sets that allowed all types in the model to be endemic in the pre-vaccination steady state.

For each included parameter set, the following steps were executed:

- simulate the pre-vaccination steady state (as defined in Section C.2) and compute predictors $HR_{VT,i}$, $HR$, $OR_{VT,i}$ and $OR$ to predict type-specific and overall replacement;
- simulate the 'true' outcome of vaccination at the post-vaccination steady state (as defined in Section C.2) and compare it to the pre-vaccination predictions.

After simulating all parameter sets, the proportion of corrected predictions is computed for type-specific and overall outcome. When there were multiple non-vaccine types, the type-specific performance was first defined and computed for each non-vaccine type separately. As the interactions between different types were generated randomly, the performance of each non-vaccine type should be identical on average. Hence, we computed the numbers shown in Figure 2 of the main text and in eFigure 5 by averaging over the performance of all non-vaccine types.

| Parameter | Values or range |
|---|---|
| Number of types ($n$) | $2, 3, \cdots, 7$ |
| Per capita contact rate ($c$) | † |
| Probability of successfully acquiring type $i$ given contact with infected ($\beta_i$) | Uniformly distributed between 2.5 and 3.5 for all type $i$† |
| Per capita clearance rate of type $i$ ($\mu_i$) | 1 for all type $i$ |
| Interaction parameters in acquisition ($k_{ij}$) | Uniformly distributed between 1/3 and 3 on a log scale |
| Interaction parameters in clearance ($h_{ij}$) | Uniformly distributed between 1/3 and 3 on a log scale |
| Parameter for asymmetric interactions ($\epsilon$) | $0.2, 0.5$ |

eTable 1: Parameter values or ranges used in simulations. † As the per capita contact rate $c$ and acquisition probabilities $\beta_i$ always appear in product. The value of $c$ is absorbed in $\beta_i$ for notational convenience, which now may attain values above one.

### F Decomposition into direct and indirect interactions

We here rewrite predictor $HR_{1,3}$ (3) into expression (9) under the pairwise-symmetric multiplicative structure with only interaction in clearance by decomposing expression (3) into direct and indirect interactions.

According to definition (3),

$$\begin{aligned}
HR_{1,3} &= \frac{I_{1|\{1,12\}}q_{1\to 13} + I_{12|\{1,12\}}q_{12\to 123}}{I_{\emptyset|\{\emptyset,2\}}q_{\emptyset\to 3} + I_{2|\{\emptyset,2\}}q_{2\to 23}} \\
&= \frac{\frac{I_1}{I_1+I_{12}}q_{1\to 13} + \frac{I_{12}}{I_1+I_{12}}q_{12\to 123}}{\frac{I_\emptyset}{I_\emptyset+I_2}q_{\emptyset\to 3} + \frac{I_2}{I_\emptyset+I_2}q_{2\to 23}} \\
&= \left(\frac{I_1 q_{1\to 13} + I_{12}q_{12\to 123}}{I_1 + I_{12}}\right) / \left(\frac{I_\emptyset q_{\emptyset\to 3} + I_2 q_{2\to 23}}{I_\emptyset + I_2}\right) \\
&= \left(\frac{I_1 k_{13} q_{\emptyset\to 3} + I_{12}k_{13}k_{23}q_{\emptyset\to 3}}{I_1 + I_{12}}\right) / \left(\frac{I_\emptyset q_{\emptyset\to 3} + I_2 k_{23} q_{\emptyset\to 3}}{I_\emptyset + I_2}\right) \\
&= k_{13} \cdot \left[\left(\frac{I_1 + I_{12}k_{23}}{I_1 + I_{12}}\right) / \left(\frac{I_\emptyset + I_2 k_{23}}{I_\emptyset + I_2}\right)\right] \\
&= k_{13} \cdot \left[\frac{(I_1 + I_{12}k_{23}) \cdot (I_\emptyset + I_2)}{(I_1 + I_{12}) \cdot (I_\emptyset + I_2 k_{23})}\right].
\end{aligned}$$

The next step is to rewrite $I_1$ and $I_{12}$ using the lack of interaction in clearance and the detailed balance property (i.e. $I_X q_{X\to Y} = I_Y q_{Y\to X}$ as proved in Section D):

$$I_1 = I_\emptyset \frac{q_{\emptyset\to 1}}{q_{1\to\emptyset}},$$

$$I_{12} = I_2 \frac{q_{2\to 12}}{q_{12\to 2}} = I_2 \frac{k_{12} q_{\emptyset\to 1}}{q_{12\to 2}} = I_2 k_{12} \frac{q_{\emptyset\to 1}}{q_{1\to\emptyset}}.$$

Using the above equalities, the equivalence of expressions (3) and (9) can be shown:

$$\begin{aligned}
HR_{1,3} &= k_{13} \cdot \left[\frac{(I_\emptyset \frac{q_{\emptyset\to 1}}{q_{1\to\emptyset}} + I_2 k_{12} \frac{q_{\emptyset\to 1}}{q_{1\to\emptyset}} k_{23}) \cdot (I_\emptyset + I_2)}{(I_\emptyset \frac{q_{\emptyset\to 1}}{q_{1\to\emptyset}} + I_2 k_{12} \frac{q_{\emptyset\to 1}}{q_{1\to\emptyset}}) \cdot (I_\emptyset + I_2 k_{23})}\right] \\
&= k_{13} \cdot \left[\frac{(I_\emptyset + I_2 k_{12} k_{23}) \cdot (I_\emptyset + I_2)}{(I_\emptyset + I_2 k_{12}) \cdot (I_\emptyset + I_2 k_{23})}\right].
\end{aligned}$$

### G Two scenarios of incorrect prediction

Here, we show two simulated scenarios of incorrect prediction in a three-type system with $VT = \{1\}$ and $NVT = \{2, 3\}$, under the pairwise-symmetric multiplicative structure and only interaction in acquisition. In eFigure 4, the steady-state prevalence is shown for increasing effective vaccine coverage. In the simulation, the effective vaccine coverage was implemented as a reduction factor that acts on the acquisition hazard of the vaccine types. In both scenarios, elimination of the vaccine type was reached at an effective vaccine coverage of around 60%.

eFigure 4a shows a scenario with $k_{12} = 0.5$, $k_{13} = 0.9$ and $k_{23} = 0.4$. The predictor for type 3 indicated replacement ($HR_{1,3} = 0.92 < 1$), while replacement did not occur. Studying the decomposition of $HR_{1,3}$ into direct and indirect interactions (see expression (9) in the main text) shows that the component for direct (indirect) interaction indicated the tendency of promoting (preventing) replacement. However, the resulting $HR_{1,3}$ predicted replacement, while the indirect interaction was actually stronger, preventing replacement.

eFigure 4b shows a scenario with $k_{12} = 1.4$, $k_{13} = 0.8$ and $k_{23} = 0.2$. The overall predictor indicated replacement $HR = HR_{1,2} \cdot HR_{1,3} = 1.46 \cdot 0.66 = 0.96 < 1$, while replacement did not occur. However, the overall non-vaccine-type prevalence decreased only slightly.

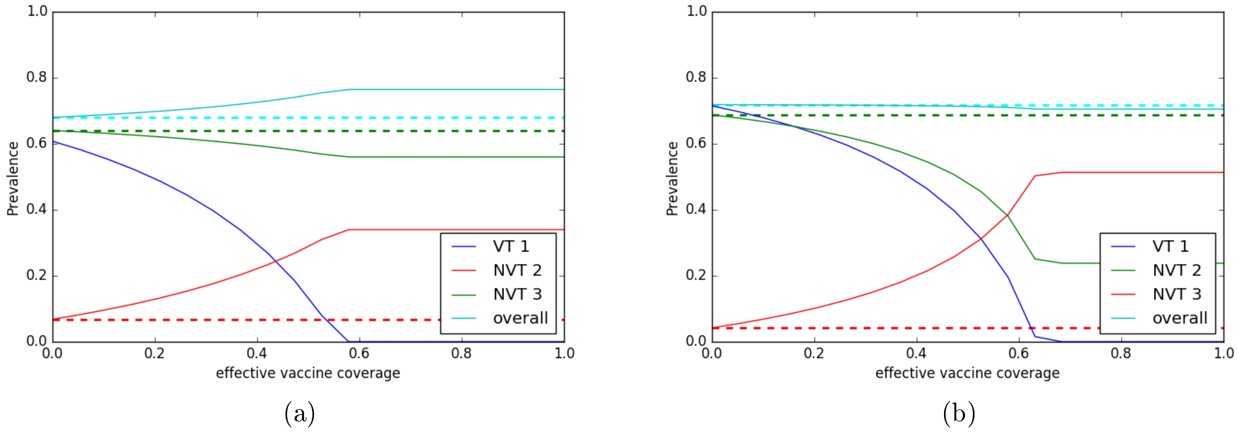


eFigure 4: Two scenarios of incorrect prediction in a three-type system with $VT = \{1\}$ and $NVT = \{2, 3\}$ under the pairwise-symmetric multiplicative structure and with only interaction in acquisition. The steady-state prevalence is shown for increasing effective vaccine coverage. The pre-vaccination prevalence is shown by dashed lines, whereas the post-vaccination prevalence is shown by solid lines at values of the effective vaccine coverage above 60%. (a) A scenario with opposing direct and indirect effects. The predictor for type 3 indicated replacement incorrectly. (b) A scenario with modest decrease in the overall non-vaccine-type prevalence, while the overall predictor indicated replacement incorrectly.

### H Simulation results with interactions in acquisition and clearance

In this section, we present the simulation results obtained from the model including interaction in clearance, in addition to that in acquisition. It was simulated following the same procedure as outlined in Section E. Each parameter set was expanded by interaction parameters for clearance, which were generated in the same manner as the corresponding interaction parameters for acquisition. For instance, the parameter sets under the pairwise-symmetric multiplicative structure each included $n(n-1)/2$ parameters $h_{ij}$ that were uniformly generated on a log scale in the interval $(1/3, 3)$ by means of Latin hypercube sampling. Note that for the groupwise-symmetric structure, the same groups were used to define interactions in acquisition and clearance.

The performance of predictors $HR_{VT,i}$, $HR$, $OR_{VT,i}$, and $OR$ under the alternative multiplicative structures for interactions are shown in eFigure 5.

Pairwise-symmetric with interaction in clearance:

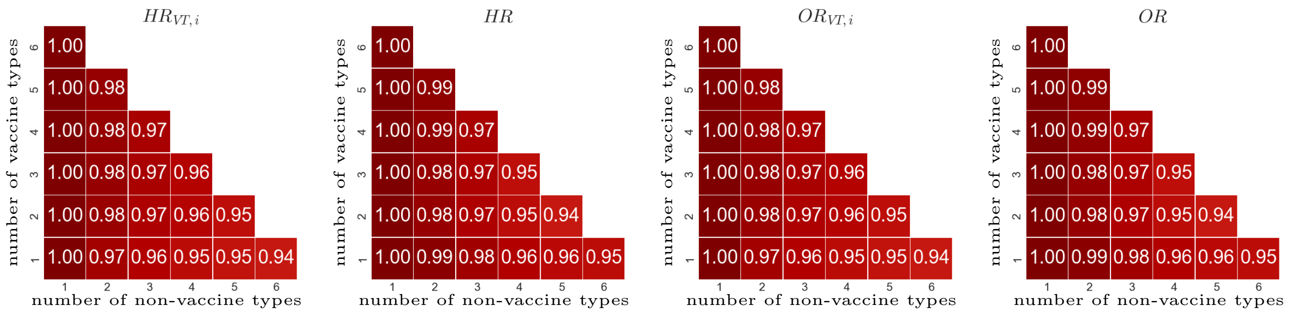


Groupwise-symmetric with interaction in clearance:

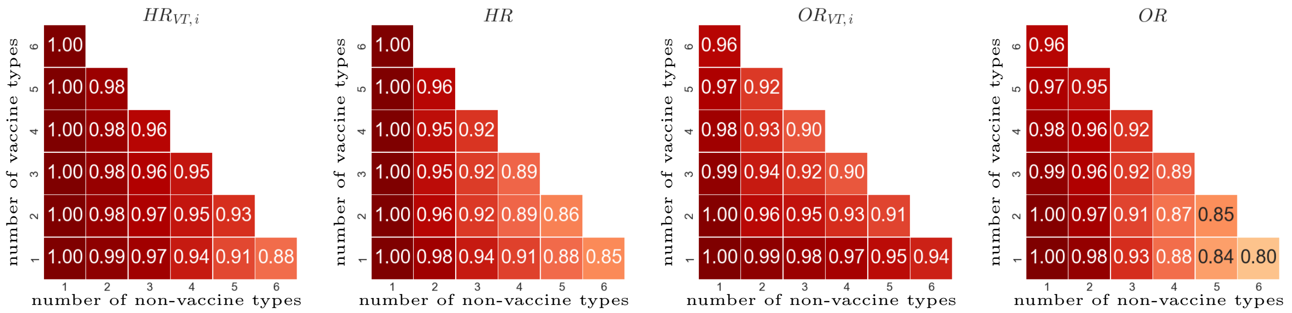


Mildly pairwise-asymmetric ($\epsilon = 0.2$) with interaction in clearance :

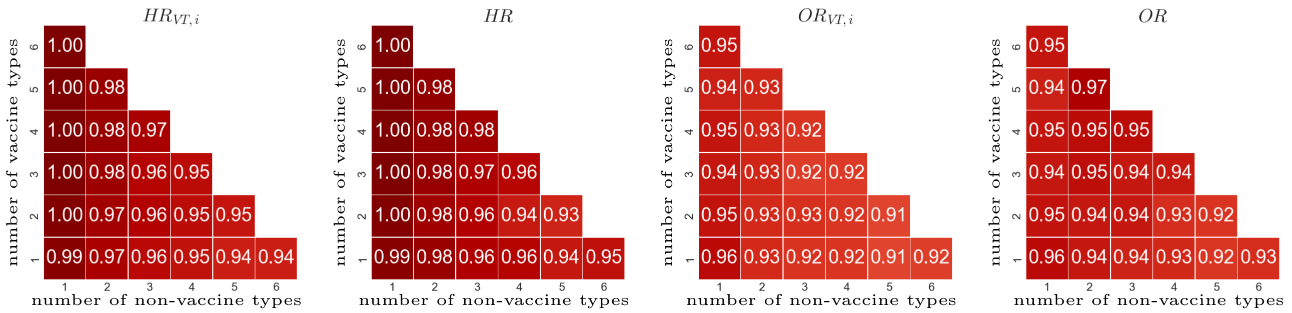


Moderately pairwise-asymmetric ($\epsilon = 0.5$) with interaction in clearance:

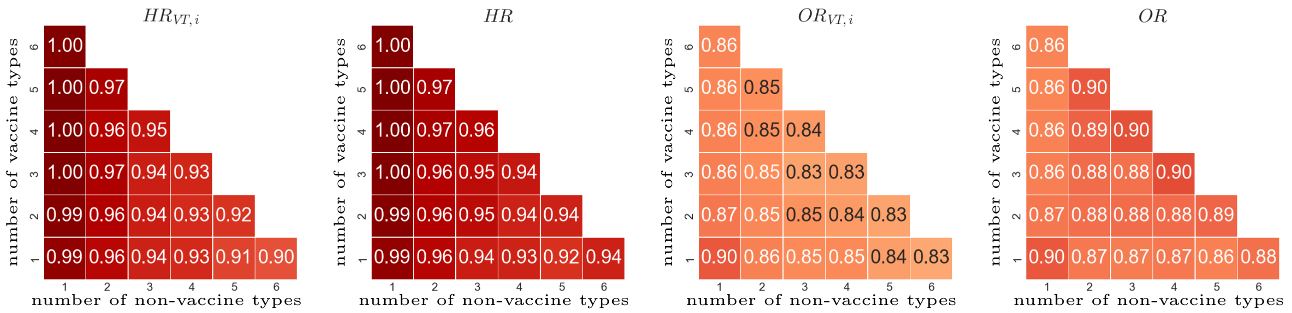


Strongly pairwise-asymmetric with interaction in clearance:

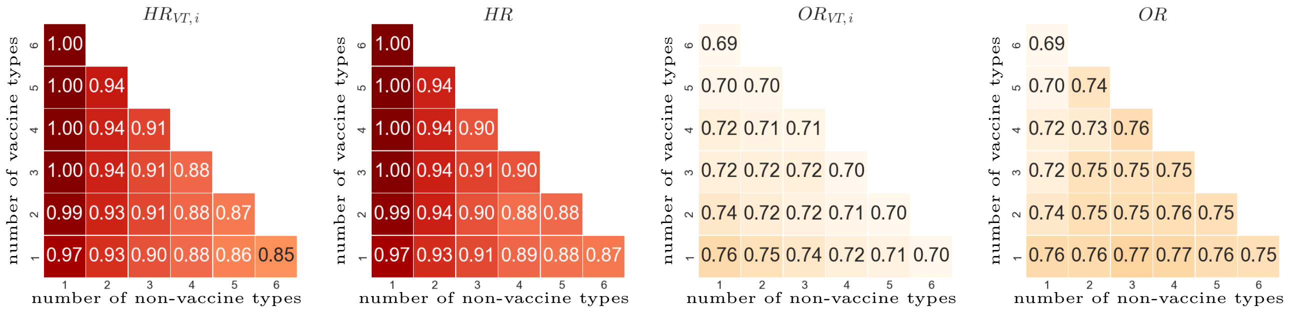


eFigure 5: Performance (proportion of correct predictions among all generated parameter sets) of predictors $HR_{VT,i}$, $HR$, $OR_{VT,i}$, and $OR$ under alternative multiplicative structures for interactions in acquisition and clearance. Row 1: pairwise-symmetric. Row 2: groupwise-symmetric. Rows 3-5: increasing pairwise-asymmetric with increasing values of $\epsilon$ for more asymmetry between pairs of reciprocal interaction parameters. Performance of $HR_{VT,i}$ and $OR_{VT,i}$ were obtained by averaging over the performance of each non-vaccine type $i$.

#### Electronic supplementary material references:

[1] Grimmett G, Stirzaker D. *Probability and random processes.* Oxford university press, 2001, 237-239.
